# Cortico-hippocampal dynamics of hierarchical syntactic planning in natural speech production

**DOI:** 10.64898/2026.08.06.743237

**Authors:** Piermatteo Morucci, Mamady Nabé, Sebastian Sauppe, Martin Meyer, Pierre Mégevand, Laurent Spinelli, Balthasar Bickel, Timothée Proix, Anne-Lise Giraud

## Abstract

The human brain must rapidly construct hierarchical structures to organize complex sequential behavior, yet the neural dynamics supporting this process during natural behavior remain poorly understood. Spoken language provides a powerful model system for investigating this computation, requiring rapid transformation of conceptual intent into structured sequential output. Using rare intracranial stereo-electroencephalography (SEEG) recordings from patients producing extended spontaneous speech, we examined how syntactic planning unfolds over time using measures of constituency, dependency structure, and probabilistic syntactic categories. We identified a hierarchical planning architecture in which global sentence structure and core syntactic categories (nouns and verbs) were specified before more local planning operations. Neural representations of these categories emerged up to 1 s before articulation and persisted throughout the planning period, whereas optional modifiers, including adjectives and adverbs, were recruited only closer to speech onset. These observations support a model of hierarchical incremental planning in which abstract sentence structure precedes the incremental specification of individual sentence elements. While core syntactic categories engaged a broader fronto-temporo-parietal network than other word classes, syntactic-depth-related activity emerged in parallel across cortical regions and the hippocampus, suggesting that hippocampal relational representations contribute to sentence structure building. Together, these findings support a cortico-hippocampal model of speech production in which hierarchical sentence structure and core syntactic categories are planned before secondary syntactic elements are incrementally incorporated into the evolving sentence plan. These results provide a neural account of how abstract linguistic structure is transformed into fluent speech.

## Introduction

Spoken language is a highly efficient and structured means of conveying information, enabling humans to rapidly transform abstract thoughts into well-formed utterances. Central to this ability is grammatical encoding, in which conceptual content is mapped onto hierarchical syntactic structures and corresponding lexical items. Within this framework, syntactic linearization refers to the process by which abstract hierarchical representations are converted into ordered word sequences for articulation^1,2^.

Despite extensive previous work on language production, the precise neural computations allowing the brain to dynamically construct syntactic structures when planning speech remain largely unresolved, particularly under naturalistic conditions. The use of artificial or highly constrained production tasks (e.g.,^3–5^) limits generalization to spontaneous speech. The few studies employing naturalistic stimuli predominantly rely on neuroimaging techniques with poor temporal resolution (e.g.,^6,7^), limiting the ability to track planning operations in real-time. This body of work has identified a distributed set of cortical regions implicated in hierarchical syntactic representations and syntactic linearization, including the left inferior frontal gyrus (LIFG), the left posterior middle temporal gyrus (LpMTG), the anterior and posterior superior temporal gyri (aSTG, pSTG), and the temporal pole^8,9^.

More recently, studies combining naturalistic paradigms with high-density intracranial recordings have implicated portions of a frontotemporal circuit centered on the caudal part of the inferior frontal cortex (pars triangularis and pars opercularis) and the caudal middle frontal gyrus as critical contributors to speech planning^10,11^. However, these studies did not isolate the planning of syntactic structure from other linguistic computations such as lexical retrieval, semantic encoding, or prosodic planning. Thus, although previous research has delineated cortical regions involved in syntactic planning, a major gap remains in understanding the temporal dynamics of syntactic structure generation during natural speech.

Planning coherent syntactic structures during natural speech entails the coordination of semantics with multiple syntactic subroutines. A first set of speech planning operations involves retrieving and assembling fundamental syntactic objects at the word level, namely syntactic categories (e.g., nouns, verbs, adjectives). Among these, core syntactic classes (e.g., nouns and verbs) serve as the structural backbone of the sentence structure, providing the argument structure and predicative framework for organizing sentence meaning. In contrast, local non-core lexical categories such as adjectives and adverbs instantiate modification relations, specifying properties of entities and events. Function words (e.g., determiners and prepositions) encode grammatical relations and hierarchical dependencies, thereby supporting the integration of lexical items into well-formed syntactic configurations. These objects form the building blocks of phrasal structure and determine the hierarchical organization of the sentence^12^. Yet, when and how elements from different syntactic classes are specified during spontaneous speech planning remains unresolved. Behavioral observations indicate that longer pauses often precede the articulation of core syntactic categories, such as nouns^13^. Similarly, eye-tracking studies show that adjectives are planned relatively late, typically after the noun has been selected^14^. Together, these findings suggest that different syntactic categories are represented within distinct temporal windows during speech planning prior to articulation. Neurophysiological evidence, however, remains scarce, partly due to the methodological difficulty of studying natural speech with time-resolved techniques.

A second set of syntactic subroutines involves planning hierarchical tree-like structures that specify how individual lexical items combine into larger units such as phrases, clauses, and sentences. These include on the one hand the planning of global syntactic scaffolds — i.e., sentence-level tree structures — whose depth and configuration determine the overall structural framework of the utterance. On the other hand, it involves local, word-level bracketing operations that organize lexical items into nested phrasal projections during linearization^15^.

Besides these constituency representations, planning syntactic structures likely engages a third set of operations devoted to the computation of relational dependencies. (e.g., filler–gap dependencies, subject–predicate relations). These relations pose specific planning challenges for core syntactic categories—especially nouns and verbs — which participate in long-distance dependencies and often form the backbone of the sentence’s argument structure. Dependencies must be maintained in working memory until they are fully linearized. Finally, probabilistic factors – such as the likelihood of a specific syntactic class or sequence – likely play an important role in shaping syntactic planning, with more common configurations facilitating planning and less probable ones imposing greater processing demands^16^. Given the substantial number of syntactic elements and operations required to generate well-formed utterances, the human brain has likely evolved an efficient planning strategy that balances early activation of manipulable syntactic objects with minimization of concurrent load. How our brains achieve this balance remains unknown. One option is *hierarchical incrementality*, whereby core syntactic categories and global syntactic scaffolds are activated earlier than local elements and maintained until all required elements are integrated^17–19^. An alternative is *linear incrementality*, in which syntactic elements are planned according to their temporal order of articulation, with neural encoding emerging in close proximity to speech onset^20,21^.

To address this gap, we leveraged intracranial stereo-electroencephalography (SEEG) recordings from patients engaged in natural speech production. We used multiple encoding models capturing distinct dimensions of syntax to probe neural activity time-locked to each word onset across cortical and subcortical sites. This approach provides millisecond-level access to the neural dynamics of speech planning, allowing us to characterize how different levels of syntax structure are constructed and coordinated in real time (Figure 1).

**Figure 1.**
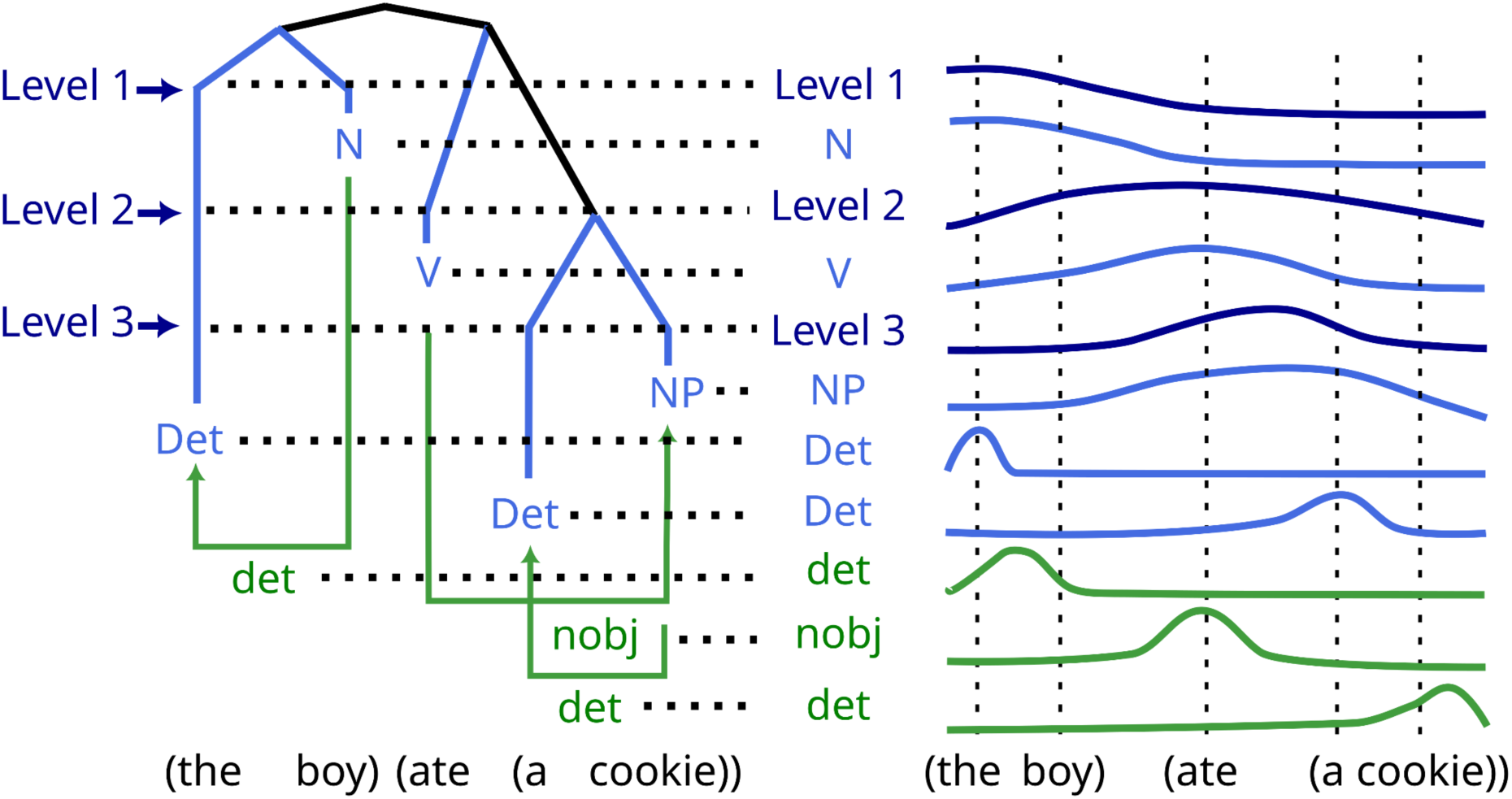
Distributed encoding of hierarchical syntactic planning in speech: Left: Constituency, dependency structure, and probabilistic lexical categories were probed in natural speech production segments. Right: intracranial recordings from patients reveals that global sentence structure and core syntactic commitments are established before and maintained longer than local lexical realization.

## Results

Stereo-electroencephalography (SEEG) was recorded in four French-speaking patients as they engaged in natural speech production tasks, involving telling a story or recalling an autobiographical episode (see *Methods*) (Figure 2, A, B). This SEEG dataset provides coverage of both cortical and hippocampal sites, allowing us to assess the contribution of these structures to syntax planning and production. We then extracted from the produced stories word-level linguistic features reflecting different aspects of the constituency, dependency and probabilistic structure of sentences. Constituency features captured representations of content-based syntactic categories, including core syntactic classes (nouns, verbs, auxiliaries) and non-core modifier classes (adjectives, adverbs), which together constitute the basic units of a constituency tree. To model global syntactic configurations, we extracted word-by-word measures of syntactic depth from constituency tree representations derived from a parser (see *Methods*), indexing each word’s position within the evolving hierarchical structure and serving as a proxy for sentence-level structural complexity. To capture local, word-level, structure-building operations, we quantified the number of openings and closings nodes at each word, reflecting incremental bracketing operations during online syntactic planning. Dependency structure was modeled by counting, at each word, the number of opening dependencies established online, capturing incremental head–dependent integration. Contextual syntactic probability features were derived using the part of speech (POS) probability distribution of next word predictions (Figure 2, B, C). These features were used to construct word-level regression models. Each model comprised a baseline set of predictors capturing variance associated with the planning of words (i.e., word onset), as well as their lexical and semantic properties (i.e., Zipf frequency and static semantic embeddings), combined with a syntactic feature of interest (e.g., syntactic depth). This modeling strategy was adopted to facilitate interpretation. We verified that the features of interest showed low intercorrelations (see supplementary figure 3).

**Figure 2.**
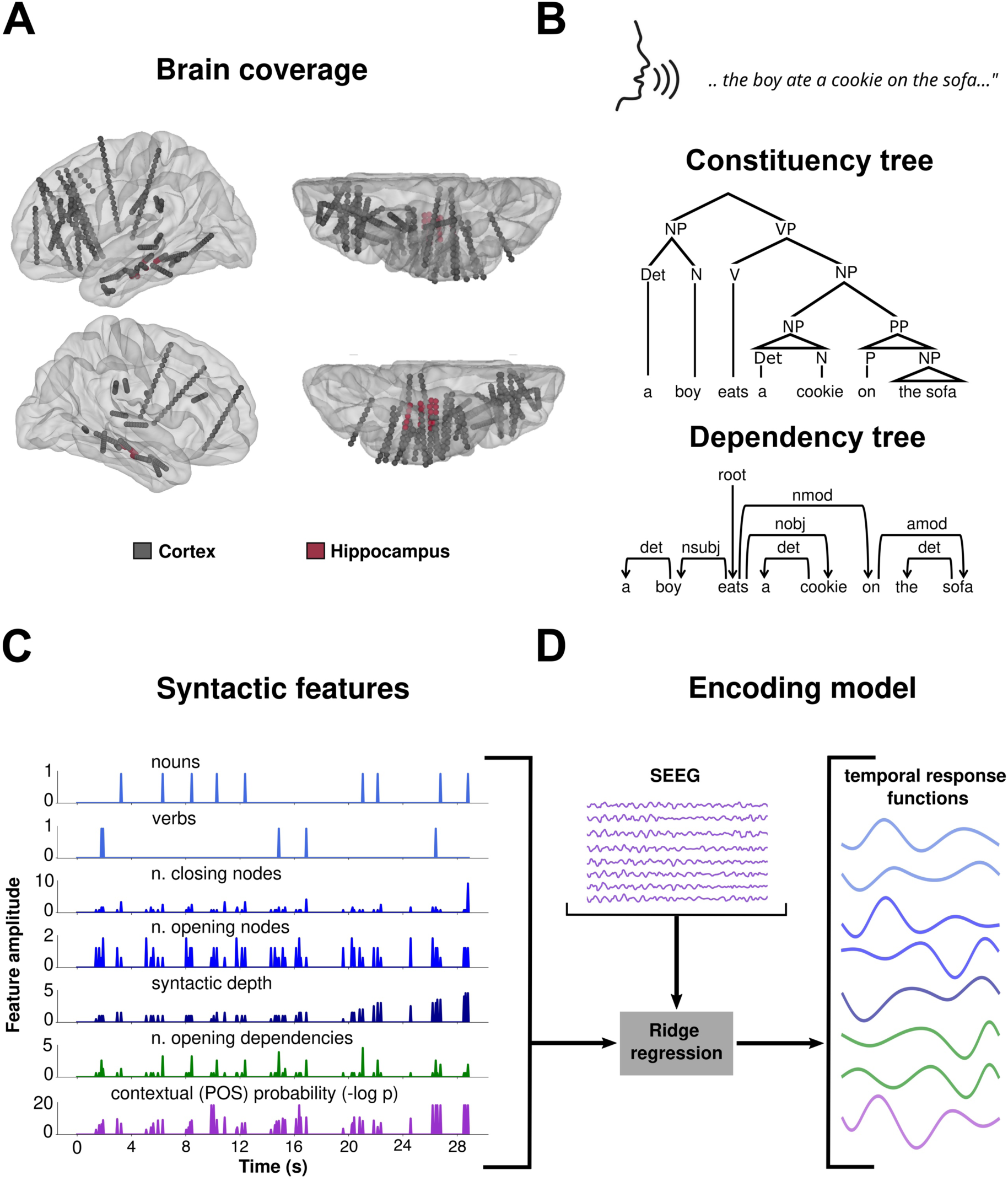
Syntactic features extracted from natural speech production data, and analysis workflow: **(A)** Distribution of neocortical and hippocampal sites from all four patients included in the study. **(B)** An example of how the same sentence can be represented either as a constituency tree or as a dependency tree, derived from their respective parsers. **(C)** Feature values of the regressors derived from the constituency and dependency syntactic representations shown in (B). Contextual Part of Speech Probability was derived independently. The plotted features refer to a selection of consecutive words produced by a patient during 28 s. **(D)** Multivariate Temporal Response Function (mTRF) encoding model probing linear relationships between features and neural signals by capturing regression weights across both sites and time lags. Regularized linear regression was applied to model the relationship between word-level syntactic features extracted from natural speech production data and the neural activity across single SEEG sites.

To assess how these syntactic features are spatially and temporally represented in neural activity, we applied a two-step analysis including first a Multivariate Temporal Response Function (mTRF)^20^ regression model using each syntactic feature as a predictor to predict neural activity at each SEEG contact (see *Methods*). We used a Laplacian montage to emphasize spatially localized neural signals. Neural activity was quantified using broadband gamma power, a well-established index of local cortical activity (see *Methods*). This approach allowed us to identify, for each feature, cortical sites showing better-than-chance accuracy in reconstructing the neural response. Second, to assess the temporal dynamics of syntactic features selection during speech planning, a cluster-based permutation test was applied to the resulting kernel of each model and contact, assessing the presence of temporal clusters^21^. Through this approach we identified, for each contact, *when* a given feature is represented in neural activity (Figure 2, C, D). In the following, we first focused on SEEG contacts located in the cortex bilaterally and later assessed the representation of syntactic structures over hippocampal sites.

### Syntactic classes

We first explored the neural representation of syntactic classes, which form the building blocks of both constituents and dependencies. We used a mTRF model that includes a binary regressor for each content word (i.e., nouns, verbs, adjectives, adverbs). We focused on content words only, as these syntactic elements are known to be more strongly represented in neural activity than function words^22^. A binary regressor for auxiliaries was also included in the model, as this syntactic class usually plays verb-like roles. Accuracy from the model was compared against a null distribution made of 100 surrogate mismatch models, and only models outperforming the null 95% of the times were considered significant (see *Methods* for details). Using this approach, we identified contacts where the syntactic class model accurately predicted the neural response, with major contributions of electrodes located in bilateral temporal regions and left frontal regions (Supplementary Figure 1; Figure 3).

**Figure 3:**
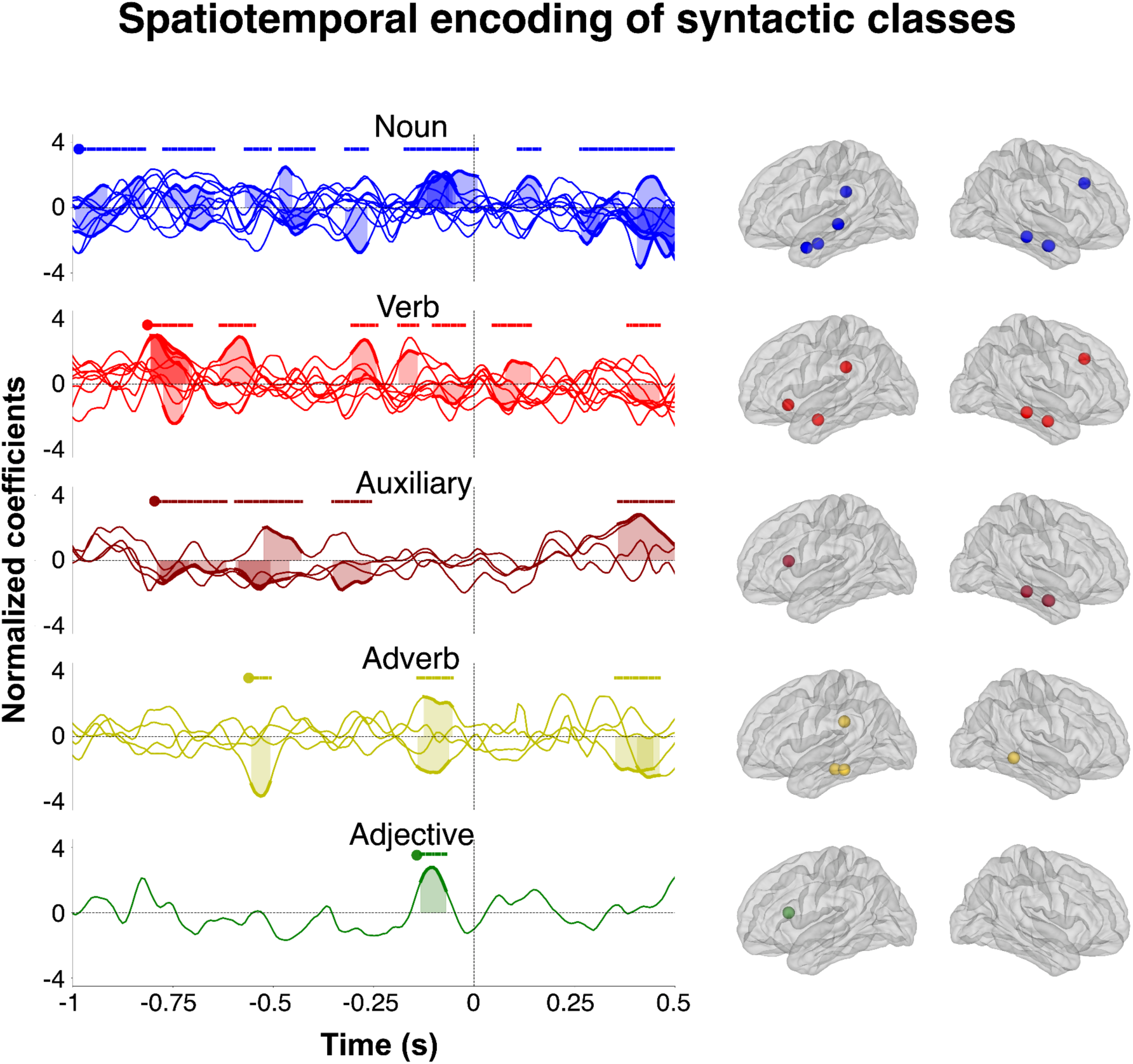
Spatiotemporal representations of syntactic classes. Time-series show the significant response-functions (kernels) for each syntactic class. The vertical dotted line pointing at time point 0 indicates word onset. The horizontal lines above the kernels indicate significant time windows. Brain plots next to the time-series show the location of the kernels projected on the brain surface of the patients. Significance was assessed for each contact individually (based on change-level prediction compared to surrogate null distribution).

To investigate whether core vs. secondary syntactic categories are differentially represented, we assessed the spatial and temporal effects associated with each syntactic element. We selected from each significant contact the response-functions (kernels) associated to syntactic classes. A cluster-based permutation test was applied across the full time window (−1 to 0.5 s; with 0 being word onset) of these kernels. Specifically, each kernel was tested against a null distribution generated through permutation to identify temporal clusters where feature-related activity significantly deviated from chance (i.e., falling outside or above the 97.5th percentile of the null distribution; see ^23^). All syntactic classes showed distinct spatiotemporal profiles. Effects related to core syntactic classes such as nouns and verbs appeared first (∼1 s and 0.8 s before articulation), and spanned several contacts over bilateral mid and inferior frontal, bilateral temporal and left parietal regions. Auxiliaries followed a similar temporal profile to verbs, with a spatial distribution restricted to right temporal and left mid-frontal regions. Adverb planning arose later, around 0.5 s before articulation, over left and right temporal contacts, while adjectives emerged just before articulation in only one contact in the left mid-frontal region. These findings show that different syntactic classes are represented in distinct spatiotemporal patterns during natural speech planning, with core classes (e.g., nouns, verbs) emerging earlier and across more widespread regions than modifiers (e.g., adjectives, adverbs). Overall, noun was the syntactic class that elicited earlier and more distributed cortical signals.

### Constituency planning

We then examined constituency planning, that is, representations reflecting hierarchical relationships between word chunks. To capture the planning of global sentence scaffolds, we first tested a model based on syntactic depth, indexing the level of each word within the syntactic tree. This model allowed us to assess whether neural activity tracked transitions across different levels of the syntactic tree. Statistical analyses parallel to that used for syntactic classes (see *Methods* for details) showed positive effects for syntactic depth across bilateral frontal, temporal and parietal regions (Figure 4, A), with dominant effects in mid frontal and mid/inferior temporal regions (Figure 4, B). Constituency planning as indexed by syntactic depth emerged early, around 1 s before articulation. Significant effects indicated a decrease in neural activity in frontal regions and an increase in temporal regions (horizontal bars in Figure 4, C).

**Figure 4:**
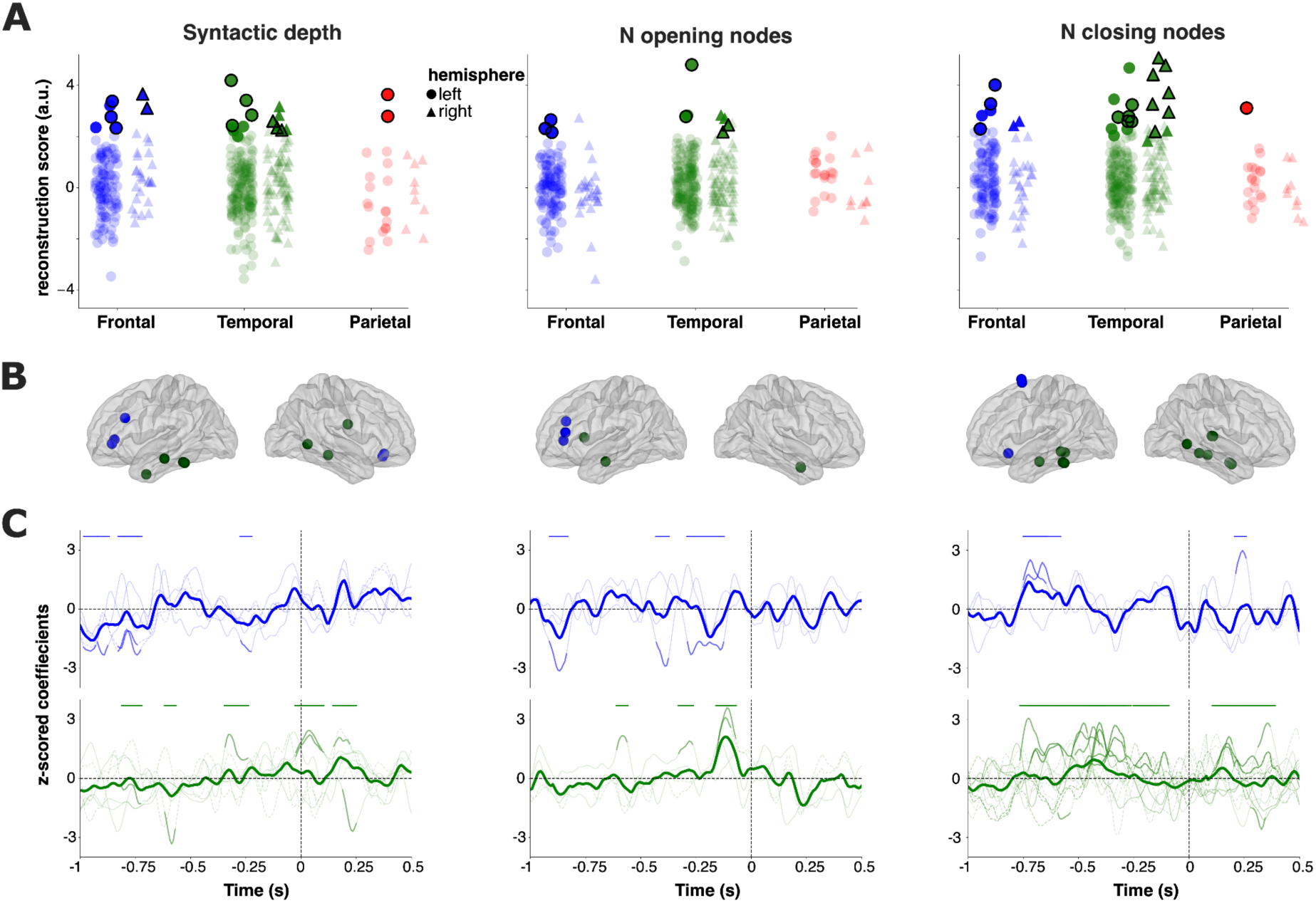
Spatiotemporal encoding of constituency features: **(A)** Accuracy of the syntactic depth, top-down and bottom-up parser model in reconstructing neural activity at each SEEG contact. Dots (left hemisphere) and triangles (right hemisphere) reflect the SEEG contacts in which accuracy is above chance. Dots and triangles outlined in black indicate the SEEG contacts in which also the kernel is significant. **(B)** Location of the cortical SEEG contacts in which the kernels are significantly projected on the patients’ mean brain surface. **(C)** Time-series showing the course of significant kernels for each constituency, over both temporal and frontal regions. The thick line shows the grand mean across significant contacts, thin lines show individual contacts. The vertical dotted line point indicates word articulation onset (time point 0). The horizontal lines show the time-windows of the kernels’ significance.

We then tested two models of local, word-level bracketing operations, which track the number of syntactic nodes opening and closing at each word. Both models relied on the same underlying tree, but differed in their node-counting procedures. The first model, which counts opening nodes at each word (also referred to as “top-down model”), emphasizes anticipatory planning of upcoming constituents. The second model, counting closing nodes (also referred to as “bottom-up”), captures the completion of constituents and the modulation of neural activity at phrase and sentence boundaries. We found significant effects of opening and closing node models over frontal and temporal contacts associated with distinct spatio-temporal profiles (Figure 4, A, B). Positive effects of the *opening nodes model* emerged slightly later than those of syntactic depth, around 0.9 s before articulation with a majority of kernels being significant over 0.4 s to 0.15 s before articulation. This effect was associated with an activity decrease in prefrontal regions and an increase in temporal ones.(Figure 4, C). This pattern suggests that these regions might work in tandem to support the planning of syntactic node openings (Figure 4, C). The *closing-node model* was associated with a stronger and more lasting effect over temporal than frontal regions, emerging 0.75 s before articulation until 0.2s, and reemerging around 0.25 s after articulation. In both regions, closing nodes were associated with neural activity increase (Figure 4, C).

### Dependency planning and contextual POS probability

In addition to the two previous analyses, we examined the spatiotemporal dynamics underlying the planning of syntactic dependencies and contextual syntactic probability. To this end, we tested (i) a model based on the number of opening syntactic dependencies derived from a dependency-tree representation of the sentences, and (ii) a purely probabilistic model based on the probability of POS based on the previous 4 words. The number of opening dependencies was primarily encoded in bilateral mid/superior frontal and left mid-temporal regions (Figure 5, A, B). This feature was characterized by sustained neural activity decrease over the 0.4 s before articulation, with only one contact in the inferior frontal sulcus showing early encoding at 0.9 s before articulation (Figure 5 C). Effects of POS probability also appeared over bilateral temporal and frontal regions, in particular over portions of the left inferior and mid-frontal cortex, and the right superior frontal cortex. The POS probability time-course indicates that this feature is primarily associated with activity decreases emerging early before articulation (∼1 s) (Figure 5, C).

**Figure 5:**
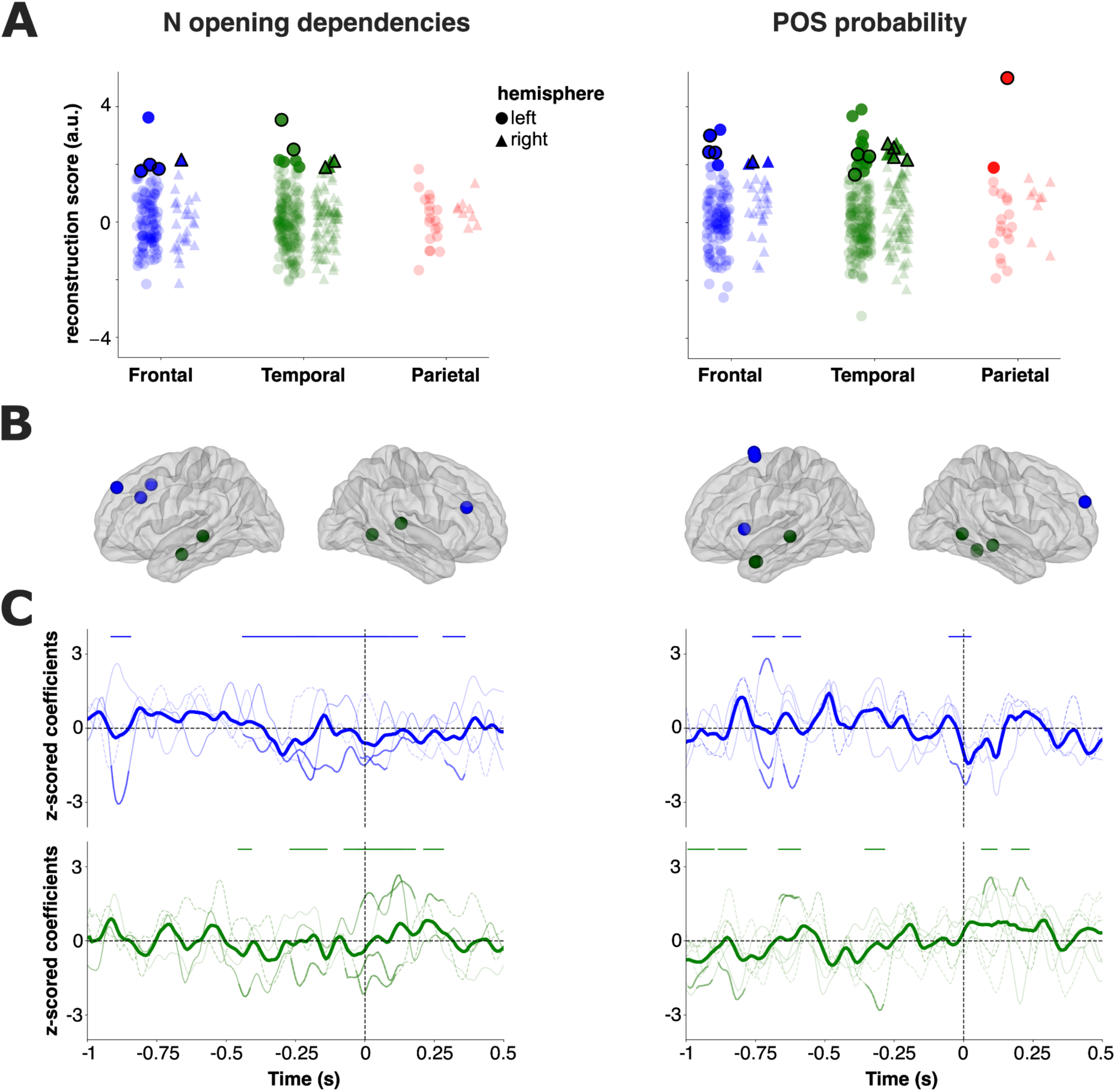
Spatio-temporal encoding of dependency relations and probabilistic features. **(A)** Accuracy of the opening dependencies and POS probability model in reconstructing neural activity at each SEEG contact. Dots (left hemisphere) and triangles (right hemisphere) reflect the SEEG contacts in which accuracy is above chance. Dots and triangles outlined in black indicate the SEEG contacts where the kernel is also significant. **(B)** Locations on the cortical SEEG contacts of the significant kernels for each constituency model. **(C)** Time-series showing the significant kernels for each constituency, over both temporal and frontal regions. The thick line shows the grand mean across significant contacts, thin lines show individual contacts.The vertical dotted line at time 0 indicates word onset. Horizontal lines above the kernels illustrate the significant time-windows.

### Syntax representations in the hippocampus

While the previous analyses were limited to neocortical sites, we next examined whether any feature of the syntactic models could account for neural activity recorded in the hippocampus^24^. All four patients had at least one electrode localized in the hippocampus (left and/or right; see Supplementary Materials for anatomical details). Syntactic depth was the feature most strongly reflected in hippocampal activity (Figure 6 A). The syntactic depth effect was fairly consistent across subjects (present in 3 out of 4 patients), whereas POS probability, syntactic class and the number of opening dependencies were seen in only one patient, and only on a few contacts (see supplementary Figure 2). The timing of the effects varied between patients and kernels, stretching from 1 s to 0.25 s before articulation (Figure 6 B).

**Figure 6:**
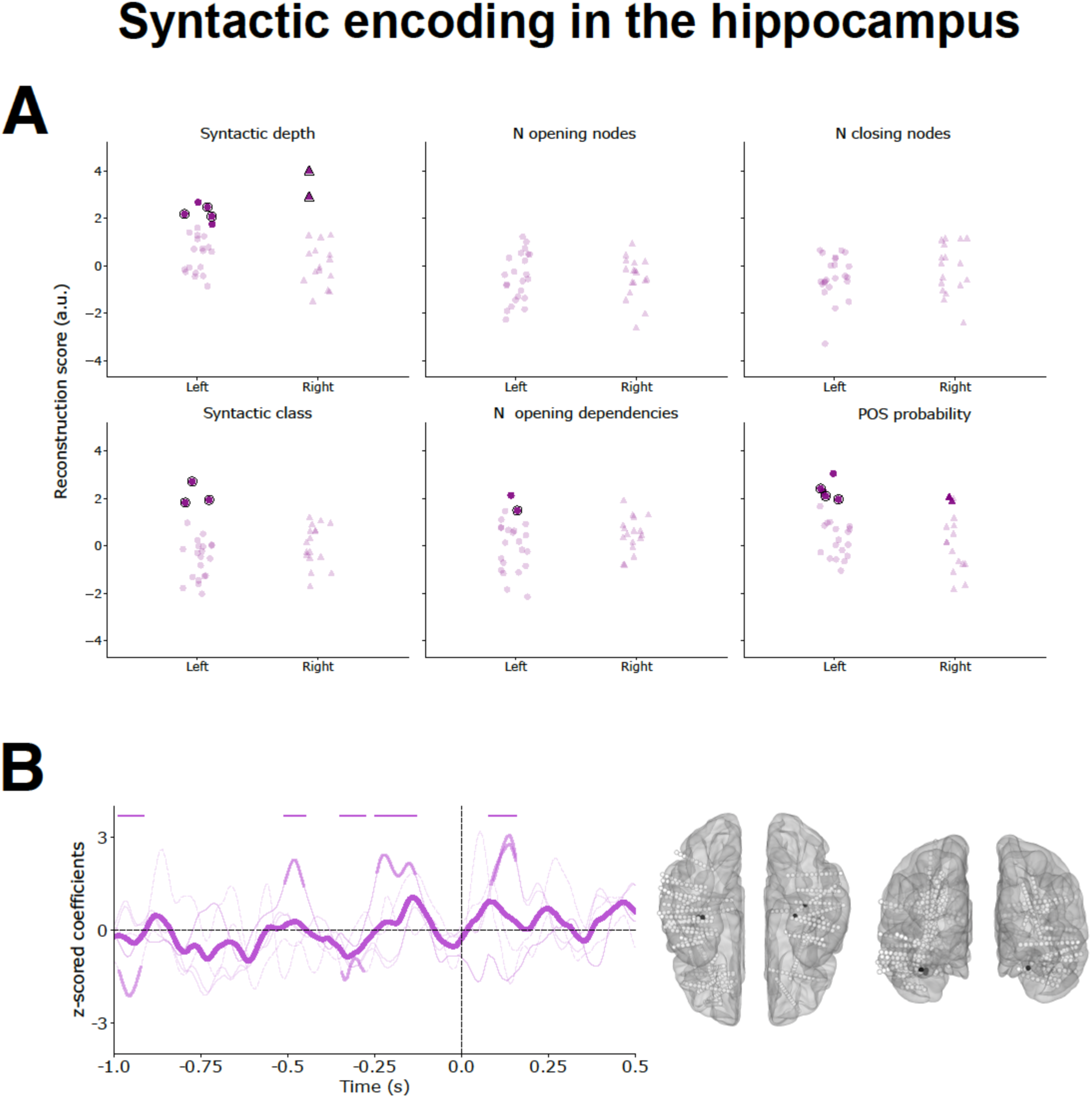
Syntax representation in the hippocampus: **(A)** Accuracy of all syntactic models in reconstructing neural activity at each SEEG contact in the hippocampus. Each dot and triangle reflect a contact in the left and right hemisphere, respectively. Coloured dots and triangles reflect the SEEG contacts where accuracy is above chance. Dots and triangles outlined circled in black indicate the SEEG contacts where the kernel is also significant. **(B)** Time-series showing significant kernels for the syntactic depth model. Vertical dotted lines at time point 0 indicate word onset. The horizontal lines above the kernels show the time-windows of the kernel’s significance. Brain plots next to the time-series show the locations of significant syntactic depth model, which is the only model fairly consistent across subjects.

## Discussion

Planning coherent sentences during natural speech production requires the coordinated integration of multiple syntactic subroutines. Using intracranial SEEG recordings from epileptic patients engaged in natural speech production, we identify a neural planning scheme that enables the brain to accomplish this task by flexibly coordinating multiple manipulable syntactic objects and structures, efficiently balancing the load of each subroutine. Our core finding indicates that syntactic planning is highly anticipatory and follows a specific deployment: the core scaffold of sentence structure, as probed at the word level by the depth level of a constituency tree and core syntactic categories, is selected about 1s before articulation, maintained throughout the planning process across frontal and temporal regions. In contrast, local linearization processes involved in bracketing operations of open and closing nodes and modifiers (e.g., adverbs, adjectives) are resolved later, in closer temporal proximity to articulation, and are associated with more transient and local neural signatures. Our study also shows that the core scaffold of sentence structure, operationalized as syntactic depth, involves in parallel the hippocampus.

Our results provide evidence that different syntactic classes are planned at distinct temporal stages prior to articulation, suggesting a structured deployment (Figure 1). Core syntactic categories such as nouns, verbs, and auxiliaries (which play verb-like roles) are recruited early, approximately 1 to 0.8 s prior to articulation, and are characterized by sustained representations over time across spatially distributed patterns in bilateral temporal and frontal regions, in line with previous literature^5^. This profile is consistent with their role in establishing the structural backbone of the sentence, including argument structure and event representation, and suggests that these elements must be maintained in working memory over an extended window to support subsequent planning operations. In contrast, modifier categories (adjectives and adverbs) are represented closer to articulation (around 0.5 s prior), and are associated with more spatially restricted and transient activity patterns. These categories often function as optional modifiers that do not usually constitute a structural commitment within the sentence global frame and can therefore be incorporated at a later stage without interfering with the overall structure. The spatio-temporal dissociation between core and modifier syntactic classes is difficult to reconcile with strictly linear incremental accounts of syntactic processing^25,26^. Instead, it supports a model in which core aspects of hierarchical structure are specified ahead of local elements selection^27^, in line with previous behavioral^17^ and ECoG studies^19^ showing evidence of early hierarchical planning.

Of particular interest is the finding that nouns, a core syntactic category in sentence structure, are planned earlier, over a longer time period, and in a more distributed manner than other syntactic classes^28^. The planning of nouns mobilizes more resources because it results from decisions not based locally but on preceding discourse, assumptions about the interlocutor’s state of knowledge (which affects the choice between lexical nouns vs pronouns vs ellipsis), the way they integrate with a verb as arguments (which affects whether they need special marking such as prepositions). This interpretation is consistent with behavioral evidence showing longer planning pauses before nouns are being pronounced^13^, suggesting they are associated with greater preparation prior to articulation. The more distributed neural pattern (in space and time) may further reflect the need to integrate lexical, semantic, pragmatic, and syntactic information across multiple systems to establish stable referential representations that can support subsequent combinatorial operations. The early activity may also reflect that the noun is a core element of the sentence that needs to be activated early irrespective of its exact lexical content.

Accordingly, core structural representations are established prior to local bracketing operations. The early emergence of syntactic depth effects, beginning up to ∼1 s before articulation across bilateral prefrontal and temporal cortices, suggests that speakers generate an abstract, tree-like scaffold of the upcoming utterance well ahead of speech onset. Local bracketing operations, indexed by opening and closing nodes, emerged later prior to articulation, with the planning of opening nodes preceding the planning of closing nodes. The temporal ordering of these effects shows a planning architecture in which a global hierarchical framework is first established and subsequently maintained, while local operations dynamically update this structure to support linearization. Interestingly, opening-node effects were characterized by opposing activity patterns in prefrontal (decreases) and temporal (increases) regions with frontal decreases preceding temporal increases. The temporal offset between frontal deactivation and temporal activation could suggest a transition between successive stages of hierarchical sentence planning, whereby frontal regions disengage following the establishment of new hierarchical planning stage (node opening), allowing temporal regions to instantiate node-specific linguistic representations. However, the functional roles of these antagonistic effects remain to be elucidated.

In contrast, closing-node effects were characterized by joint increased activity in temporal and frontal regions, mirroring effects previously reported in the comprehension literature^29^, and were present both before and after articulation. The opposing dynamics observed at node opening and closing suggest that sentence planning is organized into distinct phases of hierarchical reconfiguration and integration. While node opening is characterized by a transient decoupling between frontal and temporal activity, consistent with structural expansion and content specification. In contrast, node closing is associated with joint frontal–temporal increases, suggesting a network-wide cohesive integration process that both finalizes syntactic dependencies and consolidates the evolving sentence structure. Post articulatory effects could signal a similar sequence at sentence completion.

Examining how syntactic planning engages dependency relations and local probabilistic constraints, we show that the encoding of the number of opening dependencies has similar spatio-temporal dynamics to opening nodes, pointing to a common representational strategy for planning anticipatory commitments during sentence construction. In contrast, the earlier effects associated with POS probability are more consistent with sensitivity to local sequential expectations, such that more frequent configurations appear to facilitate planning while less probable structures impose greater processing demands. This suggests that sentence planning involves both hierarchical syntactic scaffolding and incremental, probability-based processes, operating in parallel but with different functional roles in guiding the unfolding utterance.

Our dataset included deep contacts located in the hippocampus, offering a unique window into its contribution to syntax planning. Among the several features tested, syntactic depth was the only one reliably represented in the hippocampus activity across patients, pointing to a selective involvement of this region in encoding the global backbone of sentence structure. Syntactic depth was encoded well before articulation, highlighting the contribution of the hippocampus in hierarchical syntactic representation building. In accordance with the role of the hippocampus in domain-general relational representations and predictive planning across cognitive domains^30,31^, this finding suggests that the hippocampus supports the speech planning process by encoding relational structures representing early transitions between levels of the syntactic tree. The hippocampus would thus not contribute to syntactic processing per se, but to maintaining or constructing the relational scaffold that organizes hierarchical sentence structure during planning, with this structured representation being accessed by the frontotemporal cortex for syntactic elaboration and incremental linearization into a speakable form.

This finding aligns with previous work showing that the hippocampal system supports the acquisition of language-like rules and the tracking of hierarchical sequences and the storage of semantic knowledge^32–34^: Damage to this system may result in impaired sentence-level language abilities, providing causal evidence for its importance in supporting syntactic processing^35,36^.

Our results show that some syntactic features have a sustained representation at the network level throughout the planning phase from 1 s before to articulation onset (e.g., nouns in Figure 2). However, this sustained representation does not appear at the level of single contacts, but emerges from transient, peak-like dynamics distributed across different contacts. This particularity suggests that the syntax scaffolding emerges from a distributed and continuous updating process within an evolving syntactic state, rather than as a set of static representations that remain stable over time. Such a dynamic organization is consistent with the demands of natural speech production, which requires continuous adjustment and refinement of the unfolding sentence structure. It may also reflect the large pool of semantic representations syntactic elements are selected from.

Together, these findings suggest that sentence planning unfolds as a structured cycle of hierarchical construction and stabilization. During planning, core syntactic categories (nouns and verbs) show early and sustained engagement within a distributed fronto-temporo-parietal network, consistent with their role in initiating and anchoring hierarchical structure construction, while syntactic depth indexes the relational organization maintained across successive construction stages. Hippocampal involvement may reflect the representation of relational structure supporting the incremental assembly of nested syntactic dependencies. Node opening is characterized by a transient dissociation between frontal and temporal activity, consistent with the establishment of a new syntactic position and the early specification of core lexical anchors. Node closing, in contrast, is associated with joint fronto-temporal activation, consistent with network-wide stabilization that integrates newly formed constituents into the evolving sentence structure. More broadly, these findings highlight the central role of syntax in transforming abstract conceptual representations into structured, time-ordered speech.

## Methods

### Participants

Stereo EEG (SEEG) recordings were obtained from four participants (2 female, mean age = 32 years; range = 30–35). All participants had pharmacoresistant epilepsy and underwent intracranial electrode implantation as part of their clinical epilepsy treatment. They were native French speakers with normal sensory and cognitive functions and exhibited left-hemisphere language dominance.

Of the four participants, three had SEEG electrodes primarily implanted in the left hemisphere, while one had main coverage over the right hemisphere (see supplementary Table 1), for a total of 610 contacts over cortical and hippocampal regions (see Figure 1, A). SEEG electrodes were localized using the iELVis and Voxeloc toolboxes^37–39^. Participants were recruited from Geneva University Hospitals (Switzerland). All provided informed consent, and the experiment was approved by the local ethical committee under the number 2021-00480.

### Speech production tasks

Spontaneous speech production was collected in three conditions. The participants either (1) watched a narrative visual movie without speech, or (2) browsed through short image books without texts, and then verbally described the stories or depicted the book images in as much detail as possible, or (3) described a past autobiographical event. They were allowed to speak for as long as they wished and indicated verbally when they were done. All the descriptions were done in French. Speech production data was recorded using a TASCAM DR-40 microphone, while brain activity was simultaneously recorded. Not all participants completed all tasks, and the duration of their speech recordings varied between 8 and 25.5 minutes, with an average of 17.03 minutes.

### Speech transcription

Audio files were transcribed using an automatic speech-to-text translator (WhisperX^40^), which returns annotations, and aligned at the word level using the Montreal Forced Aligner^41^. The transcriptions were subsequently reviewed by a native French speaker for text accuracy, quality and timing. We then manually defined sentence boundaries. Unlike written text, where sentence boundaries are clear, natural spoken language oftentimes lacks objective and self-evident delimitations. As a general rule, we consider as a sentence any coherent string of words containing at least one verb, or multiple verbs in cases of coordination or relative clauses. Since speech production is often marked by pauses, errors, hesitations, and self-corrections, these features were also taken into account when determining sentence offsets. We followed guidelines for segmenting natural speech production (e.g.,^6^). In particular, sentence boundaries were defined not only on syntactic grounds but also on prosodic and temporal criteria, including pause duration and speech rhythm. For example, coordinated clauses were treated flexibly: they were classified as a single sentence when produced with fluent delivery and minimal pause, but segmented into separate sentences when accompanied by a sufficiently long pause or a clear prosodic break indicating discourse-level separation. Overall, subjects produced a mean of 143.7 sentences (range: 72 – 289) and 2015 words (range = 1031 – 3032) (see supplementary table 1).

### SEEG processing and neural features

The raw SEEG signals were amplified and digitized at 2048 Hz, and then stored for offline analysis (Brain Quick LTM, Micromed, S.p.A., Mogliano Veneto, Italy). Subsequent preprocessing was performed with MNE-Python (3.6) following standard guidelines^38^. Continuous SEEG data from different tasks and patients were extracted and saved as separate files, each of which was preprocessed independently using the same pipeline. Noisy contacts were manually identified and removed. We then applied a low-pass and high-pass filter at 250 Hz and 0.1 Hz, respectively. A notch filter was applied to eliminate 50 Hz electrical noise and its harmonics from the signal. A Laplacian reference was then used along the shaft of each electrode to minimize volume conduction effects in the signal, with bipolar montage for the most proximal and distal electrodes. For every contact, we derived broadband high-frequency activity (BHA) in the 70–150 Hz range. This signal provides an approximate measure of the local average spiking activity of nearby neuronal populations^42,43^. To obtain it, we computed the mean amplitude across eight Gaussian band-pass filters, whose center frequencies were spaced logarithmically and whose bandwidths increased in a semi-logarithmic manner. The resulting BHA traces were then normalized (z-scored) per channel and resampled at 125 Hz.

### SEEG and Audio Alignment

The SEEG and audio signals for the four patients were temporally aligned using modality-specific synchronization procedures. For three patients, audio was recorded using a dedicated MP3 recorder, while SEEG signals were acquired with a clinical Micromed amplifier. In these cases, synchronization was achieved using a custom-made external device operating in standalone mode, which injected an identical, uniquely encoded pulse train into both the SEEG and audio recording channels every few seconds. These pulse trains served as shared temporal markers; all instances were subsequently detected and decoded in each modality, enabling the identification of corresponding trigger events and the derivation of a precise temporal mapping between SEEG and audio streams, with a jitter of up to 15 ms, which is short compared to the time windows at the word level we are considering. For the fourth patient, audio was obtained from the clinical video recording system, which is natively synchronized with the SEEG acquisition by the clinical software. In this case, alignment relied directly on the synchronization information provided by the clinical system, without the use of external triggers.

### Modeled features

All analyses were based on linear modeling. Specifically, we estimated forward encoding models that relate stimulus features to SEEG signals. These models are commonly referred to as multivariate temporal response functions (mTRFs)^20^. This approach allows assessing whether local field potential activity varies as a function of features, assuming the endogenously generated landmarks (e.g. word onset) globally follow the same TRFs patterns as external stimuli. Because in this study we are interested in assessing the neural implementation of syntactic structures, we derived word-level features that capture key properties of the underlying syntactic structure. First, we used syntactic parsers (see below for details) to extract the constituency and dependency trees for each sentence produced by the patient (Figure 1, B). These trees provide formal representations of sentence structure. Constituency trees capture how words group together into hierarchical phrases (e.g., noun phrases, verb phrases), while dependency trees represent the syntactic relations between individual words (e.g., subject–verb, verb–object). Both formalisms offer complementary perspectives on syntax: constituency emphasizes hierarchical phrase structure, whereas dependency highlights head-dependent relations (Figure 1, B).

We also extracted variables reflecting lexical and semantic word properties, which were used for creating a base model. In the following sections, we provide details on the implementation of the encoding model.

## Basis models

A set of baseline variables was included in the regression model as covariates to account for variability associated to semantic and lexical word-level properties and to provide a reference model against which the contribution of the syntactic variables of interest could be evaluated. The variables included in the baseline model are listed below.

### Word onset

this measure is a binary variable (0 or 1) indicating whether a given time point reflects or not the phonetic onset of a given word (here meant as a token). This measure was included to account for neural responses associated with the motor planning for articulation, independently of syntactic, semantic, or lexical properties.

### Word Frequency

Word frequency reflects how common a word is in a language, independently of context. It was computed using the Python library wordfreq and expressed on the Zipf scale^44^, a logarithmic measure where higher values indicate more frequent words. This measure was included to capture neural responses related to the lexical frequency of planned words.

### Semantic embeddings

Static word vectors for each word were obtained from pre-trained models^45^ (https://huggingface.co/facebook/fasttext-fr-vectors). Principal component analysis (PCA) was used to reduce the dimensionality of the original 300-dimensional space into principal components. We arbitrarily retained the first five principal components (explaining 0.07, 0.03, 0.02, 0.02, 0.02 of variance) to improve computational efficiency and simplify the regression model, while preserving some of the informative structure of the embeddings. This measure was inserted to account for activity associated with the semantic properties of planned words, independently of the sentence context during planning.

## Constituency trees

To assess whether and how the brain generates hierarchical tree-like constituency structures, constituency trees for each sentence produced by the patients were obtained using MTGPY, a transition-based parser using Flaubert as a backbone (https://gitlab.com/mcoavoux/mtgpy). The pre-trained model is a discontinuity-capable parser, trained on the union of FTB-train, Sequoia, French Question Bank (https://filesender.renater.fr/?s=download&token=782fbbd0-21c8-4e48-9f2d-2ce08483192d). Using the phrasal tree structures obtained from this parser, we derived two metrics reflecting two possible encoding schemes for planning of syntactic configurations, as well as a metric reflecting transition in the tree structure.

### N Opening Syntactic Nodes

This measure represents a “top-down” anticipatory planning scheme and is computed by counting the number of opening brackets associated with each word in a sentence. This planning scheme hypothesizes that syntactic encoding happens at the onset of phrases and sentences, and that the dimensionality of the syntactic structure decreases with the unfolding of the spoken sentence.

### N Closing Syntactic Nodes

This measure represents a “bottom-up” incremental planning scheme and is computed by counting the number of closing “brackets” associated with each word. This planning scheme hypothesizes that sentences are planned roughly on a “word-by-word” (or phrase-by-phrase) basis, with syntactic structures and sentence boundaries being planned mostly on the fly during production.

### Syntactic depth

this measure represents the level where a given word appears within a syntactic tree. A word that is deeply embedded within nested structures, i.e. lower in the tree hierarchy, has a higher value for this feature.

### Syntactic Class

this regressor coded the syntactic class of each word, restricted to content words only — nouns, verbs, adjectives, adverbs, and auxiliaries, the latter included given their verb-like functional role. This measure tests whether distinct neural activity profiles accompany the planning of different syntactic categories.

## Dependency relations

To assess when and how the brain encodes dependency relations, dependency trees for each sentence have been calculated using the HOPS parser (https://github.com/hopsparser/hopsparser/), a dependency parser using Flaubert as a pretrained model. The parsing model is trained on Sequoia. In order to measure the memory burden required to plan syntactic dependencies at each word, the following measures were calculated:

### Number of dependencies

this measure reflects the number of dependency relations that each word has in a given sentence. Because we did not have predictions about the directionality of dependency relations, we included both left and right hand dependencies.

## Contextual probabilities

### POS contextual probabilities

- left context: This variable measures how unlikely a specific word’s POS (i.e. syntactic class, such as noun, verb, adjective, etc) is, based on the preceding four words.

#### mTRF estimation

To relate the speech features to the neural data, an encoding mTRFs was used. Kernel time spans were chosen in the time range between −1 and 0.5 s from the onset of the word to account for a large possible time span of potential effects. For each patient, we ran one mTRF for each contact and each feature of interest. The model for the features of interest was constructed by including the features from the base model (i.e., Zipf frequency, word onset, static semantic embeddings) plus the feature of interest (e.g., syntactic depth). We selected this approach as syntactic features were overall not correlated with each other (supplementary figure 3), with the exception of verb and number of opening dependencies (r = 0.63), thus minimizing collinearity confounds. Estimation was carried out using ridge regression within a nested cross-validation framework (see below). Model performance was assessed by calculating Pearson’s correlation between predicted and actual neural data.

#### Cross-validation

A cross-validation approach was employed to assess the neural encoding of syntactic features. In the outer cross-validation loop, the continuous data was split into two folds: 80% for training and 20% for testing. This was iteratively repeated 5 times, resulting in 5 outer cross-validation loops. Within each outer training fold, a 5-fold inner cross-validation loop was conducted to tune the ridge regression hyperparameter, with alpha values ranging from 10⁻⁵ a 10⁸.

The optimal alpha value was then used to retrain the model on the 80% training data from the outer fold and to test it on the remaining 20%. Accuracy in this outer test fold was again assessed using Pearson correlation. This process was repeated across the five outer folds, resulting in five correlation values, from which we report the average. This procedure was carried out separately for each SEEG contact.

#### Statistical significance

To assess the statistical significance of each model and channel, we generated a distribution of 100 surrogate models by randomly shuffling the target feature across words while keeping the baseline features unchanged. This approach ensured that the baseline features were appropriately regressed to the neural data, enabling partial isolation of the relationship between the feature of interest (e.g., syntactic depth) and the neural signal. For surrogate models, we did not perform cross-validation to optimize the regularization parameter, but instead used the alpha value that yielded the best accuracy in the original model. A model was considered statistically significant if its performance exceeded the 95th percentile of the surrogate distribution.

#### Kernel null distributions

Once the contacts in which each model significantly predicted neural activity were identified, we assessed kernel-by-kernel the timing of feature encoding. To obtain kernels for each feature and contacts, we trained a new model on the full continuous data using the median alpha that returned the best model performance across the five outer folds. This was done for all contacts in which a given feature significantly predicted brain activity. A kernel null distribution was then calculated by running the same model on the full data, but with the value of the target feature randomly shuffled, using the same shuffling procedure used to calculate a null distribution. A cluster-based approach looking for temporal clusters was then used to assess which time points of the original kernels fall outside or above the 97.5th percentile, respectively of the surrogate distribution^23^.

## Acknowledgments.

This work was funded by Fondation pour l’Audition (FPA IDA11, A.G.), NCCR snf agreement #51NF40_180888 (A.G., B.B., M.M.), ANR - France 2030 - IHU reConnect (A.G.), and SNSF career grant 193542 and 225979 (T.P.).

## Competing interests

The authors declare no competing interests.

## Supplementary Figures

**Supplementary Figure 1:**
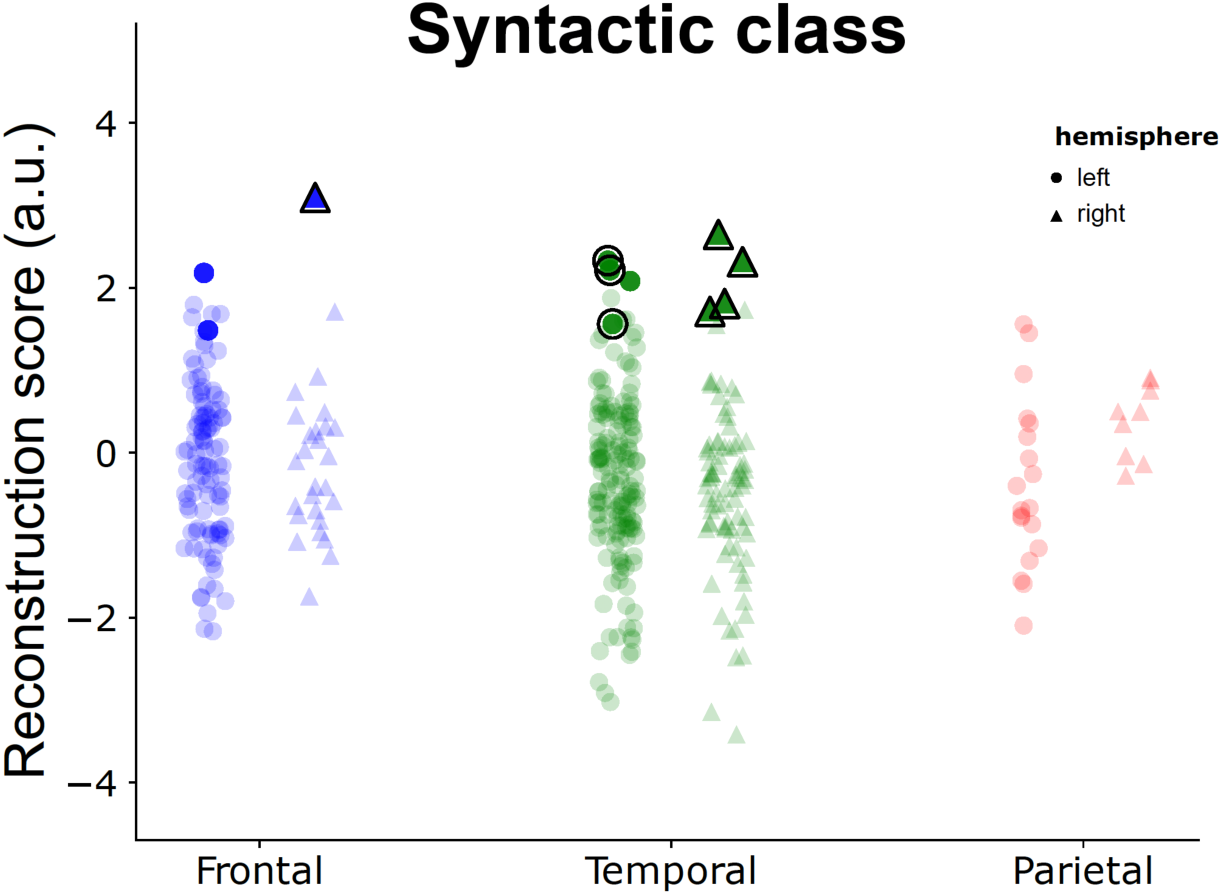
Cortical encoding of syntactic class. This figure shows the accuracy of the syntactic class model in reconstructing neural activity at each SEEG contact. Dots and triangles correspond to contacts in the left and right hemisphere, respectively. Coloured dots and triangles reflect the SEEG contacts where accuracy is above chance. Dots and triangles outlined in black indicate the SEEG contacts where also the kernels are significant.

**Supplementary figure 2:**
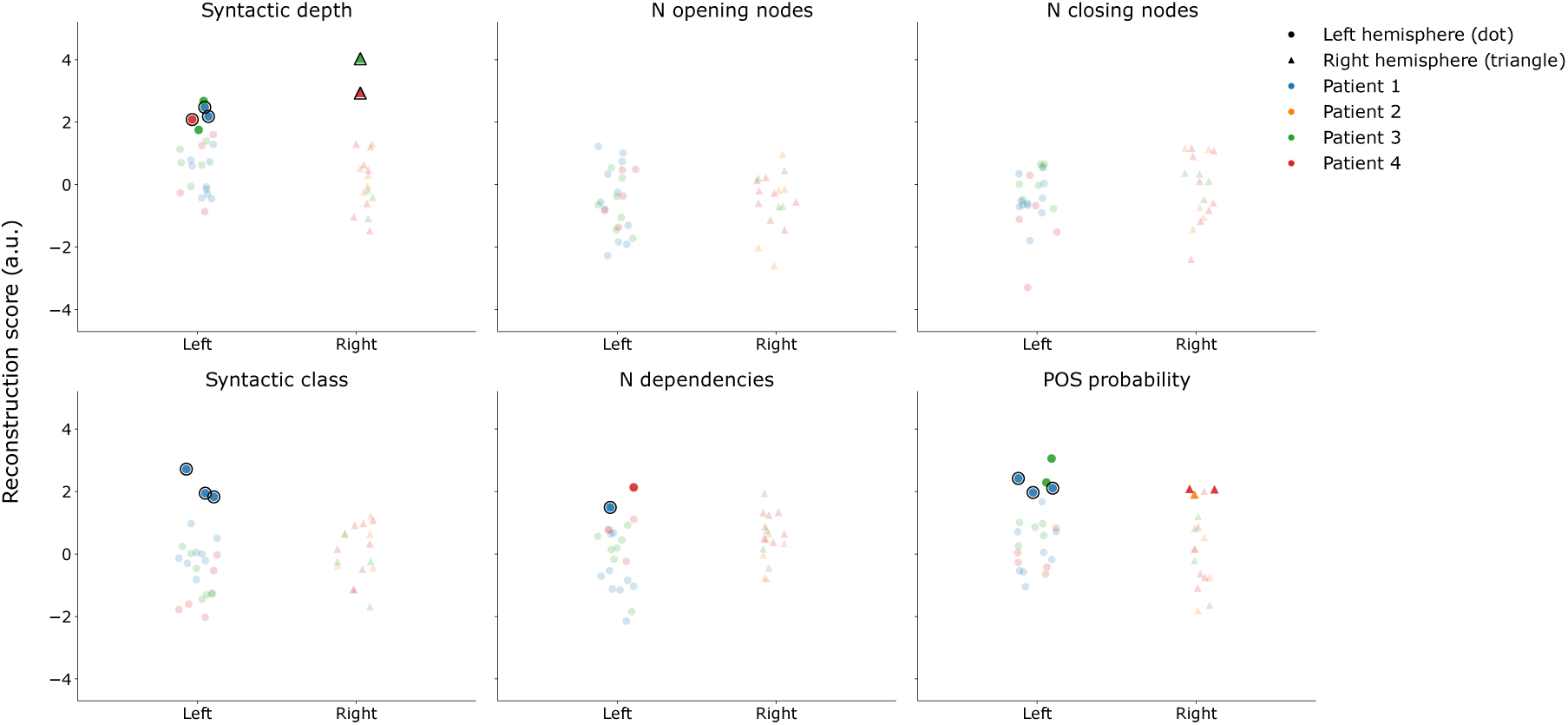
Syntactic encoding in the hippocampus: Patient-specific accuracy of all the syntactic models in reconstructing neural activity at each SEEG contact in the hippocampus. Dot and triangle correspond to contacts on the left and right hemisphere, respectively. Coloured dots and triangles reflect the SEEG contacts where accuracy is above chance. Dots and triangles outlined in black indicate the SEEG contacts in which the kernel is also significant. The color of the dot or triangle indicates the patient the electrode belongs to.

**Supplementary figure 3:**
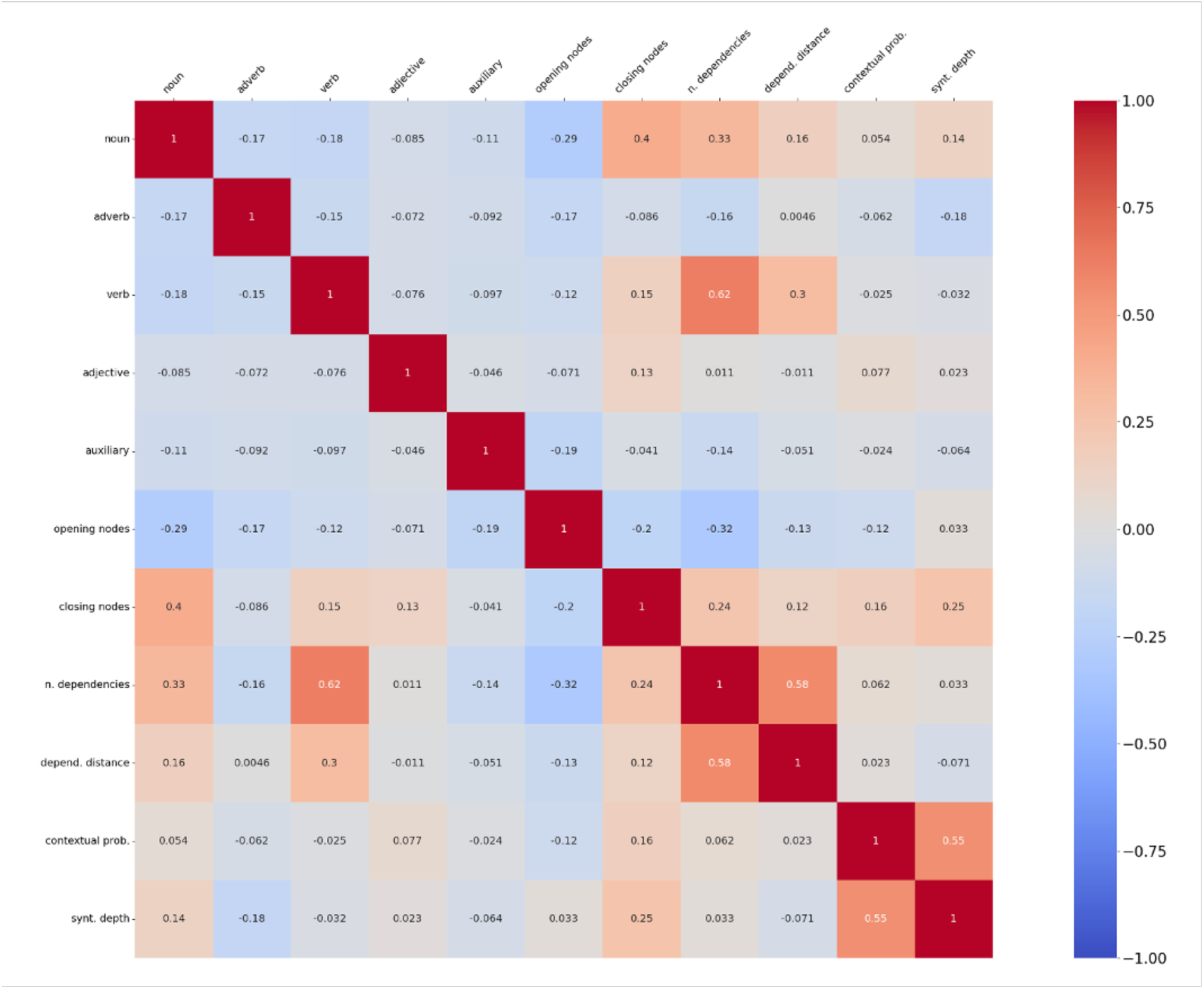
Correlation matrix of syntactic features at the word level, averaged across all patients. Colors indicate the strength and direction of pairwise correlations between features. White numbers denote statistically significant correlations.

## Supplementary tables

**Supplementary table 1:** Summary of the number of left- and right-hemisphere electrodes, the number of sentences produced, and the total number of words produced for each participant.

|  | Left hemisphere electrodes | Right hemisphere electrodes | N. of words | N. of Sentences |
| --- | --- | --- | --- | --- |
| Patient 1 | 118 | 0 | 1031 | 72 |
| Patient 2 | 166 | 10 | 3032 | 289 |
| Patient 3 | 131 | 69 | 2194 | 140 |
| Patient 4 | 0 | 116 | 1803 | 74 |

## Notes

### Competing Interest Statement

The authors have declared no competing interest.

